# Localization of the signal of dystonia-associated protein torsinA near the Golgi apparatus in cultured central neurons

**DOI:** 10.1101/2019.12.11.872804

**Authors:** Sadahiro Iwabuchi, Hiroyuki Kawano, N. Charles Harata

## Abstract

A single in-frame deletion of a codon for a glutamic acid residue within the *TOR1A* gene is linked to the autosomal-dominant movement disorder DYT1 dystonia, a condition characterized by involuntary muscle contractions that cause abnormal posture. This gene encodes the protein torsinA, and the functions of both wild-type and mutant (ΔE-torsinA) forms remain poorly understood. Previous studies based on overexpression systems indicated that wild-type torsinA resides mainly in the endoplasmic reticulum but that ΔE-torsinA is localized to the nuclear envelope or intracellular inclusions. This mutation-associated mis-localization has been proposed to underlie at least a part of the pathophysiology of DYT1 dystonia. However, the subcellular localization of torsinA has not been extensively studied when expressed at the endogenous level. Here we report an immunocytochemical analysis of torsinA proteins in cultured mouse neurons from a ΔE-torsinA knock-in model of DYT1 dystonia, where torsinA proteins are not upregulated. In all examined neurons of wild-type, heterozygous and homozygous mice, torsinA signal was found mainly near the Golgi apparatus, and only weakly in the endoplasmic reticulum and nuclear envelope. These results suggest that, in the absence of overexpression, torsinA proteins are localized near the Golgi apparatus and may influence cellular function involving the organelle.

## INTRODUCTION

DYT1 dystonia is defined as an “early-onset generalized isolated dystonia” (Albanese et al., 2013), and characterized by involuntary muscle contractions and abnormal posture. This disease is inherited in an autosomal-dominant manner, and is caused by an in-frame deletion of a single glutamic acid codon in the *TOR1A* gene (c.904_906delGAG/c.907_909delGAG; p.Glu302del/p.Glu303del), which encodes the protein torsinA (Ozelius et al., 1997). The mutant form is denoted ΔE-torsinA. Two clinical features of DYT1 dystonia are noteworthy. Firstly, neurodegeneration is not apparent (Hedreen et al., 1988; Rostasy et al., 2003; Standaert, 2011). Secondly, carriers of the mutation show enhanced excitability within the brain (Ikoma et al., 1996; Berardelli et al., 1998; Edwards et al., 2003; Defazio et al., 2007; Carbon et al., 2010a; Carbon et al., 2010b). Based on these features, the primary abnormalities are thought to be changes in neuronal functions, the synaptic microcircuitry and/or the neuronal network (Cookson and Clarimon, 2005; Breakefield et al., 2008; Tanabe et al., 2009; Niethammer et al., 2011; Quartarone and Pisani, 2011; Prudente et al., 2014). TorsinA is a protein of 332 amino acids (Ozelius et al., 1997) and belongs to the AAA+ (<u>A</u>TPases <u>a</u>ssociated with various cellular <u>a</u>ctivities) family of proteins, which generally contribute to diverse cellular processes including the trafficking, unfolding of proteins during their degradation, and the disassembly of protein complexes (Hanson and Whiteheart, 2005). Nevertheless, the details of how the ΔE-torsinA mutation leads to the symptoms of DYT1 dystonia remain unclear.

In considering the pathophysiology of DYT1 dystonia, one of the critical questions has been the subcellular localization of wild-type (WT) and ΔE-torsinA proteins within neurons (Granata et al., 2009). A definitive answer would provide insight into how the ΔE-torsinA mutation leads to neuronal dysfunction. According to current models, the WT-torsinA protein is localized mainly in the endoplasmic reticulum (ER) and also present at lower levels in the contiguous nuclear envelope. In contrast, ΔE-torsinA is preferentially located in the nuclear envelope, whose ultrastructure becomes compromised in this context, and in cytoplasmic inclusion bodies. This mutation-associated mis-localization has been hypothesized to constitute an important pathogenic event in DYT1 dystonia. However, this notion is based mainly on work with experimental systems in which human torsinA proteins are exogenously introduced and overexpressed, for instance in cultured non-neuronal cells (Kustedjo et al., 2000; Goodchild and Dauer, 2004; Naismith et al., 2004), cultured neural tumor cells (Hewett et al., 2000; Gonzalez-Alegre and Paulson, 2004; Misbahuddin et al., 2005), and cultured mouse cerebral cortical neurons (Nery et al., 2011).

It is not clear, however, whether the ΔE-torsinA is mis-localized in the native state, i.e. without overexpression, because the results of overexpression studies may not be applicable to endogenous proteins for at least several reasons. First, in carriers of the DYT1 mutation (heterozygous for the ΔE-torsinA allele, *TOR1A*^+/ΔE^), total expression of the torsinA protein is not upregulated; rather it is slightly lower than in control subjects (Goodchild et al., 2005). Second, the localization of overexpressed torsinA is affected by the degree of overexpression (Bragg et al., 2004a; Goodchild and Dauer, 2004; Naismith et al., 2004). This suggests that overexpression itself influences the localization of torsinA, and therefore could mask endogenous-level localization. Third, overexpressed torsinA proteins can induce cellular abnormalities. Specifically, the overexpression of WT-torsinA in the neurons of WT mice (*Tor1a*^+/+^) leads to an abnormality in the nuclear envelope (Grundmann et al., 2007; Kim et al., 2010), similar to one observed when a WT allele is completely absent, i.e., in homozygous torsinA knock-out mice (*Tor1a*^−/−^) and homozygous ΔE-torsinA knock-in mice (*Tor1a*^ΔE/ΔE^) (Goodchild et al., 2005; Kim et al., 2010). Fourth, heterozygous ΔE-torsinA knock-in mice (*Tor1a*^+/ΔE^), whose genotype copies that of DYT1 dystonia patients, do not have this nuclear envelope abnormality (Goodchild et al., 2005; Kim et al., 2010). Lastly, torsinA localization has not been convincingly established in a system where this protein is expressed at endogenous levels (Harata, 2014).

Here we examined the subcellular distribution of torsinA in differentiated neurons with and without torsinA overexpression. For this purpose, we used the mouse ΔE-torsinA knock-in model, which incorporates the same disease-associated mutation within the endogenous mouse torsinA gene (Dang et al., 2005; Goodchild et al., 2005; Mitchell et al., 2019). WT mice (*Tor1a*^+/+^, WT alleles only) and homozygous-mutant mice (*Tor1a*^ΔE/ΔE^, mutant alleles only) were analyzed to simplify evaluation of the localization of WT and ΔE-torsinA proteins. The heterozygous mice (*Tor1a*^+/ΔE^, one WT allele and one mutant allele) were expected to exhibit both types of localization. Levels of the torsinA protein were slightly reduced in the brains, livers and fibroblasts of the mutant vs. WT mice (Goodchild et al., 2005; Yokoi et al., 2013) even though the mRNA levels were the same among all three genotypes (Goodchild et al., 2005). Thus the torsinA proteins are not overexpressed in the ΔE-torsinA knock-in mice, and in this respect, they closely resemble fibroblasts of human patients with DYT1 dystonia (Goodchild and Dauer, 2004; Goodchild et al., 2005; Hewett et al., 2008). We examined torsinA localization in primary cultures of central neurons obtained from the cerebral cortex, striatum and hippocampus, prepared as reported previously (Kakazu et al., 2012a; Kakazu et al., 2012b; Iwabuchi et al., 2013a; Iwabuchi et al., 2013b; Koh et al., 2013; Koh et al., 2015). The use of such cultures enabled us to examine the neurons of homozygous mice, which normally do not survive to adulthood (Dang et al., 2005; Goodchild et al., 2005; Tanabe et al., 2012). It also allowed for high-resolution spatial imaging, with minimal stray noise from adjacent tissues.

Although our analysis of overexpressed torsinA proteins reproduced previous findings on torsinA localization to the ER, nuclear envelope and cytoplasmic inclusion bodies, our immunocytochemical experiments showed that the signals of endogenous torsinA proteins in the absence of overexpression were preferentially localized near the Golgi apparatus. Moreover, the signal localization was indistinguishable among neurons of different genotypes, and among different rodent species. Contrary to previous findings, these results indicate that endogenous torsinA proteins are associated with the Golgi apparatus irrespective of whether they are of the WT or mutated form. The current study indicates that existing models for DYT1 dystonia pathophysiology might need reappraisal.

## MATERIALS AND METHODS

### Genotyping and animals

All animal care and procedures were approved by the University of Iowa Animal Care and Use Committee, and were performed in accordance with the standards set by the National Institutes of Health Guide for the Care and Use of Laboratory Animals (NIH Publications No. 80-23, revised 1996). Every effort was made to minimize suffering of the animals. On postnatal days 0-1, pups of the ΔE-torsinA knock-in mouse model (Goodchild et al., 2005), of both sexes, were genotyped using a fast-genotyping procedure (EZ Fast Tissue/Tail PCR Genotyping Kit, EZ BioResearch LLC, St, Louis, MO) (Kakazu et al., 2012a). Newborn rat pups of both sexes were obtained from Crl:CD(SD) (Sprague-Dawley) rat mothers (Charles River Laboratories International, Wilmington, MA).

### Neuronal culture

Neurons from individual newborn mouse pups were cultured separately by a method described previously (Koh et al., 2013; Koh et al., 2015). In brief, the cerebral cortex, striatum and hippocampus were dissected at postnatal days 0-1. Cerebral cortex was taken from the region immediately dorsal to the striatum (Martella et al., 2009), which includes the motor cortex (Hall and Lindholm, 1974). The hippocampus included the CA3-CA1 region, but excluded the dentate gyrus (Harata et al., 2006; Kakazu et al., 2012a). The dissected tissues were trypsinized and mechanically dissociated. The cells were plated on 12-mm coverslips (thickness No. 0, Carolina Biological Supply, Burlington, NC) previously seeded with a glial feeder layer prepared from rat hippocampus, in 24-well plates. Cells from the cerebral cortex, striatum and hippocampus were plated at a density of 10,000, 24,000 and 12,000 cells per well, respectively. The cells were cultured in a humidified incubator at 37°C, with 5% CO_2_. Rat hippocampal neurons were isolated and cultured using the same method (Koh et al., 2015), and plated at a density of 6,000 cells per well. The cultured neurons were analyzed after 14-19 days *in vitro* (DIV). For each genotype, the experimental data were obtained from 3 separate culture batches (i.e., pups).

### Immunocytochemistry

Our published protocol (Koh et al., 2013; Mitchell et al., 2018) was used with slight modifications. The cultured neurons were fixed with 4% paraformaldehyde (15710, Electron Microscopy Sciences, Hatfield, PA) and 4% sucrose in Tyrode’s solution (containing, in mM: 125 NaCl, 2 KCl, 2 CaCl_2_, 2 MgCl_2_, 30 D-glucose, 25 HEPES, ∼310 mOsm without adjustment; pH 7.4, adjusted with 5 M NaOH) for 30 min at 4°C. The neurons were washed twice with Tyrode’s solution (5 min per wash) at 4°C. They were permeabilized with 0.1% Triton X-100 in Tyrode’s solution for 10 min, and washed with Tyrode’s solution three times (5 min per wash). They were blocked with 2% normal goat serum (G-9023, Sigma-Aldrich, St. Louis, MO) in phosphate-buffered saline (PBS, 70011-044, pH 7.4, GIBCO-Life Technologies) (blocking solution), for 60 min at room temperature. Thereafter they were treated with the following primary antibodies (diluted in blocking solution) overnight (15-21 hours) at 4°C.

Rabbit polyclonal antibodies used were: anti-torsinA (ab34540, Abcam, Cambridge, MA; 400x dilution), and anti-glucose-regulated protein of 78 kDa / binding immunoglobulin protein (GRP78/BiP) (PA1-014A, Thermo Fisher Scientific, Waltham, MA; 4000x dilution). Mouse monoclonal antibodies were: anti-Golgi matrix protein of 130 kDa (GM130) (610822, BD Biosciences, San Jose, CA; 400x dilution), anti-protein disulfide isomerase (PDI) (ADI-SPA-891, Enzo Life Sciences / StressGen, Farmingdale, NY; 400x dilution), and anti-nuclear pore complex (NPC) proteins (ab24609, Abcam; 400x dilution).

Following washing with PBS 3 times (7 min per wash), the neurons were incubated with goat anti-rabbit IgG antibody conjugated with Alexa Fluor 488 dye (A-11070, Life Technologies; 1000x dilution in blocking solution), and anti-mouse IgG antibody conjugated with Alexa Fluor 568 dye (A-11019, Life Technologies; 1000x dilution in blocking solution) for 60 min at room temperature. They were washed with PBS at least 5 times (20 min per wash), transferred to an imaging chamber (RC-26, Warner Instruments, Hamden, CT), and observed directly in PBS.

In some experiments, the nuclei were counterstained using membrane-permeant Hoechst 33342 (Life Technologies). After the final wash in the immunocytochemical procedure, the neurons were treated with the dye at 0.5 µg/ml in PBS for 10 min at room temperature, and washed for 3-5 min with PBS. The neurons were then transferred to the imaging chamber.

### TorsinA overexpression

Cultured rat and mouse hippocampal neurons were transfected 6-11 days after plating on coverslips, with a plasmid construct encoding either the human WT- or ΔE-torsinA tagged with GFP (Naismith et al., 2004), using the calcium phosphate-based CalPhos Mammalian Transfection Kit (631312, Clontech Laboratories, Mountain View, CA) (Harata et al., 2006). Briefly, transfection was carried out in 24-well culture plates, using 2 μg DNA per well in Minimum Essential Medium (MEM) (51200-038, GIBCO-Life Technologies, Grand Island, NY). Cells were exposed to the precipitate at 37°C for 40-70 min, washed three times with MEM, and returned to the original culture medium. In a modified transfection protocol, the DNA and buffer were mixed to prepare precipitate-forming solution in 8 small-volume steps (Jiang and Chen, 2006), rather than in a single step (Harata et al., 2006), to improve the efficiency of transfection (Jiang and Chen, 2006). Transfected cells were observed on ∼18 DIV, live and without fixation.

### Cell imaging

Imaging was confined to cultures that formed confluent or nearly confluent glial layer underneath neurons, as judged by differential interference contrast (DIC) microscopy or phase-contrast microscopy. The selected neurons did not display signs of deterioration, such as the formation of large, spherical intracellular vacuoles, beaded dendrites, clustered somata, or bundled neurites. For quantitative analysis, imaging was restricted to neurons whose proximal dendrites or somata did not overlap with those of neighboring cells, because those regions contain Golgi apparatus (Mitchell et al., 2018). All imaging experiments were performed at room temperature (23-25°C).

### Fluorescence imaging system

For general characterization by widefield optics, the cells were imaged using an inverted microscope (Eclipse-TiE, Nikon, Melville, NY) equipped with an interline CCD camera (Clara, Andor Technology, Belfast, UK). The camera was cooled at −45°C by an internal fan. Alexa Fluor 488 dye was excited using a 490-nm light-emitting diode (LED, CoolLED-Custom Interconnect, Hampshire, UK) and imaged with a filter cube (490/20-nm excitation, 510-nm dichroic long-pass, 530/40-nm emission). Alexa Fluor 568 dye was excited using a 595-nm LED (CoolLED-Custom Interconnect) and imaged with a filter cube (590/55-nm excitation, 625-nm dichroic long-pass, 665/65-nm emission). The LED was used at 100% intensity, and exposure time was 1 sec. Hoechst 33342 was excited using a 400-nm LED (CoolLED-Custom Interconnect) at 100% intensity, and imaged with a filter cube (405/40-nm excitation, 440-nm dichroic long-pass, 470/40-nm emission) at a 0.2-sec exposure. 16-bit images were acquired using a 40x objective lens (Plan Fluor, numerical aperture 1.30, Nikon) with a coupler (0.7x) and without binning, in the single-image capture mode of the Solis software (Andor). Focus levels were recorded by the Perfect Focus System (Nikon) as the absolute distance (vertical height) from the coverslip-water interface.

For quantitative characterization, the cells were imaged using a confocal laser-scanning microscope (510 confocal, Carl Zeiss MicroImaging GmbH, Göttingen, Germany) equipped with an inverted microscope (Axiovert 100M, Carl Zeiss) and a 40x objective lens (Plan-Neofluor, numerical aperture 1.30). Alexa Fluor 488 dye was excited using the 488-nm line of an argon laser at 1% intensity, and imaged using a 505-530 nm band-pass emission filter. Alexa Fluor 568 dye was excited using the 543-nm line of a helium-neon laser at 100% intensity, and imaged using a 585 nm long-pass emission filter. Voxel sizes were 0.44 x 0.44 x 0.40 µm or 0.22 x 0.22 x 0.40 µm (x-y-z). Pinhole size was 58 µm with an Airy unit of 0.8. Pixel time was 2.51 µs. Four frames were averaged to give a single 12-bit image at each level of focus (optical section), and a stack of images (z-stack) was acquired at focus levels 0.40-µm apart. Vertical height was measured as a multiple of the vertical step increase between two focus levels.

Imaging conditions were the same for each brain region, so that the absolute intensities in different cells could be compared using the same imaging modality (widefield and confocal optics).

### Image analysis

Images were opened in 16-bit format and analyzed using the ImageJ software (W. S. Rasband, National Institutes of Health, Bethesda, MD).

For quantification of the intensity of torsinA staining, the effect of noise was minimized by averaging three consecutive images in a confocal z-stack and filtering with the Gaussian-blur function. The radius of Gaussian filter was 1.0 µm with pixel size of 0.44 x 0.44 µm, or 0.5 µm with pixel size of 0.22 x 0.22 µm. For measuring the torsinA intensity in the Golgi apparatus, regions of interest (ROI’s) were assigned using the GM130 channel, with the intensity threshold that had been set as the mean intensity + 3x standard deviations in a negative control image (primary antibody omission) acquired under the same conditions. For measuring torsinA intensity in the ER, ROI’s were manually assigned in the PDI channel. For measuring intensity in the nuclear envelope and nucleoplasm, a thresholded image of the NPC was used to assign a ring-shaped ROI representing the nuclear envelope, and its interior region was assigned as the ROI for the nucleoplasm. The ER region excluded the Golgi apparatus, and the nucleoplasm excluded the nucleoli. These ROI’s were transferred to the torsinA channel and the pixel intensities therein were averaged.

In both widefield and confocal images, a background intensity was measured in a non-cellular region, at a focus level above the soma (z-direction) and in a region distant from neurons (x-y direction). This value was subtracted from all measurements of individual neurons.

For the purpose of representing intensities in all sections of a neuron, the images in a confocal z-stack were projected to a single image using the maximum intensity projection (MIP) algorithm. This algorithm projects the 3-dimensional structure to a 2-dimensional image by identifying the maximal intensity of a single pixel at all focal levels (i.e. maximal intensity at different z coordinates for a single x-y coordinate pair).

For purpose of presentation, the images were filtered using the Gaussian-blur function. They were then converted from 16-bit to 8-bit format, with the minimal and maximal intensity in individual images assigned values of 0 and 255, respectively, except where otherwise specified. Where intensity was compared under the same imaging conditions, the intensity range was determined using the whole series of images for comparison, and converted using the same intensity scale.

### Characterization of antibodies used in the current study

The anti-torsinA antibody used is a rabbit polyclonal antibody. It was used for immunohistochemical staining of the brains of WT mice (Puglisi et al., 2013), the brains of transgenic mice overexpressing ΔE-torsinA (Napolitano et al., 2010; Sciamanna et al., 2011), and cultured brain neurons of ΔE-torsinA knock-in mice (Koh et al., 2013). It was also used in Western blotting experiments to detect total torsinA protein (Cao et al., 2010; Kim et al., 2010; Yokoi et al., 2010; Gordon et al., 2011; Puglisi et al., 2013; Yokoi et al., 2013). As is the case for other anti-torsinA antibodies (Goodchild et al., 2005), it detects both WT- and ΔE-torsinA proteins (Cao et al., 2010; Napolitano et al., 2010; Sciamanna et al., 2011; Koh et al., 2013). The specificity of this antibody was demonstrated in negative controls, by immunohistochemical staining of the following samples: brains of conditional *Tor1a* knock-out mice (Liang et al., 2014); brains of WT mice following antigen preadsorption (Puglisi et al., 2013); and cultured rodent brain cells following antigen preadsorption (Koh et al., 2013). The negative control also includes the Western blotting of brain homogenates of *Tor1a* knock-out mice (Yokoi et al., 2010).

Anti-GM130 is a mouse monoclonal antibody that recognizes a *cis*-Golgi marker. The immunocytochemical labeling pattern in the current study was the same as that observed in cultured rat hippocampal neurons (Horton and Ehlers, 2003; Horton et al., 2005), and mouse cerebral cortex neurons (Vitry et al., 2010), as well as cultured astrocytes isolated from the mouse cerebral cortex (Ramamoorthy and Whim, 2008) and cultured human adenocarcinoma line HeLa cells (Roy et al., 2012; Simpson et al., 2012). It was also observed in neurons *in vivo*, in the rat hippocampus (Horton et al., 2005; Cox and Racca, 2013), the rat cerebral cortex (Horton et al., 2005), the mouse hippocampus (Clarke et al., 2009) and the mouse cerebral cortex (Condon et al., 2013).

Anti-PDI is a mouse monoclonal antibody that recognizes an ER marker. This antibody was used for immunocytochemical staining of human brains (Walker et al., 2002a), HeLa cells (Chadrin et al., 2010) and cultured mouse fibroblasts (Kornak et al., 2001).

Anti-GRP78/BiP is a rabbit polyclonal antibody that recognizes an ER marker. This antibody has been used for immunocytochemical staining of rat brain gliosarcoma 9L cells, where it co-localized with PDI (Sun et al., 2006), as well as of the human retinal pigment epithelium (Marmorstein et al., 2000).

Anti-NPC is a mouse monoclonal antibody that recognizes a nuclear envelope marker. This antibody detects nucleoporins according to the vendor information. This antibody has been used for immunocytochemical staining of the mouse hippocampal line HT22 cells (Benz et al., 2010), HeLa cells (Umlauf et al., 2013), and Chinese hamster ovary CHO cells (Chen et al., 2011).

As a control for secondary antibodies (Saper, 2005), the primary antibodies were omitted from the immunocytochemical procedures in each experiment. The imaging conditions and contrast of acquired images were the same as for the other samples, and no labeling was observed (Koh et al., 2013).

### Drugs

All chemical reagents were purchased from Sigma-Aldrich unless otherwise specified.

### Statistical analysis

Means were compared using the unpaired Student’s t test. All significance values provided are two-tailed p values.

## RESULTS

### Subcellular distribution of overexpressed torsinA proteins in rodent neurons

We examined whether we can reproduce the results of overexpression studies in previous reports. For this purpose, we transfected cultured rodent brain neurons with constructs encoding GFP-tagged WT- or ΔE-torsinA using a conventional calcium-phosphate method, and monitored GFP fluorescence. In the dendrites of hippocampal neurons (focus level as judged by DIC, 0 μm height), GFP signal from the WT construct was diffuse (arrows, Fig. 1A). Several μm higher in the z direction, at the level of soma, the fluorescence remained diffuse outside the nucleus (arrow). The signal of overexpressed ΔE-torsinA-GFP was barely detectable in dendrites and appeared diffuse at the bottom of soma (0 μm, Fig. 1B). At the somatic level, the fluorescence was localized around the nucleus in a ring (perinuclear pattern, arrow). A plot of intensity along the line drawn through the soma (line-intensity analysis, Fig. 1 bottom panels) underscores the differences in the WT- and ΔE-torsinA distributions. These distributions are consistent with localization in the ER and nuclear envelope, respectively, as reported for cultured cerebral cortical neurons of WT mice (Nery et al., 2011). Notably within individual neurons, the GFP signal showed different distribution patterns across focal levels, i.e., a signal that was local and discrete at one focal level could appear diffuse at another. Thus, caution is required in acquiring and interpreting data.

**Fig. 1.**
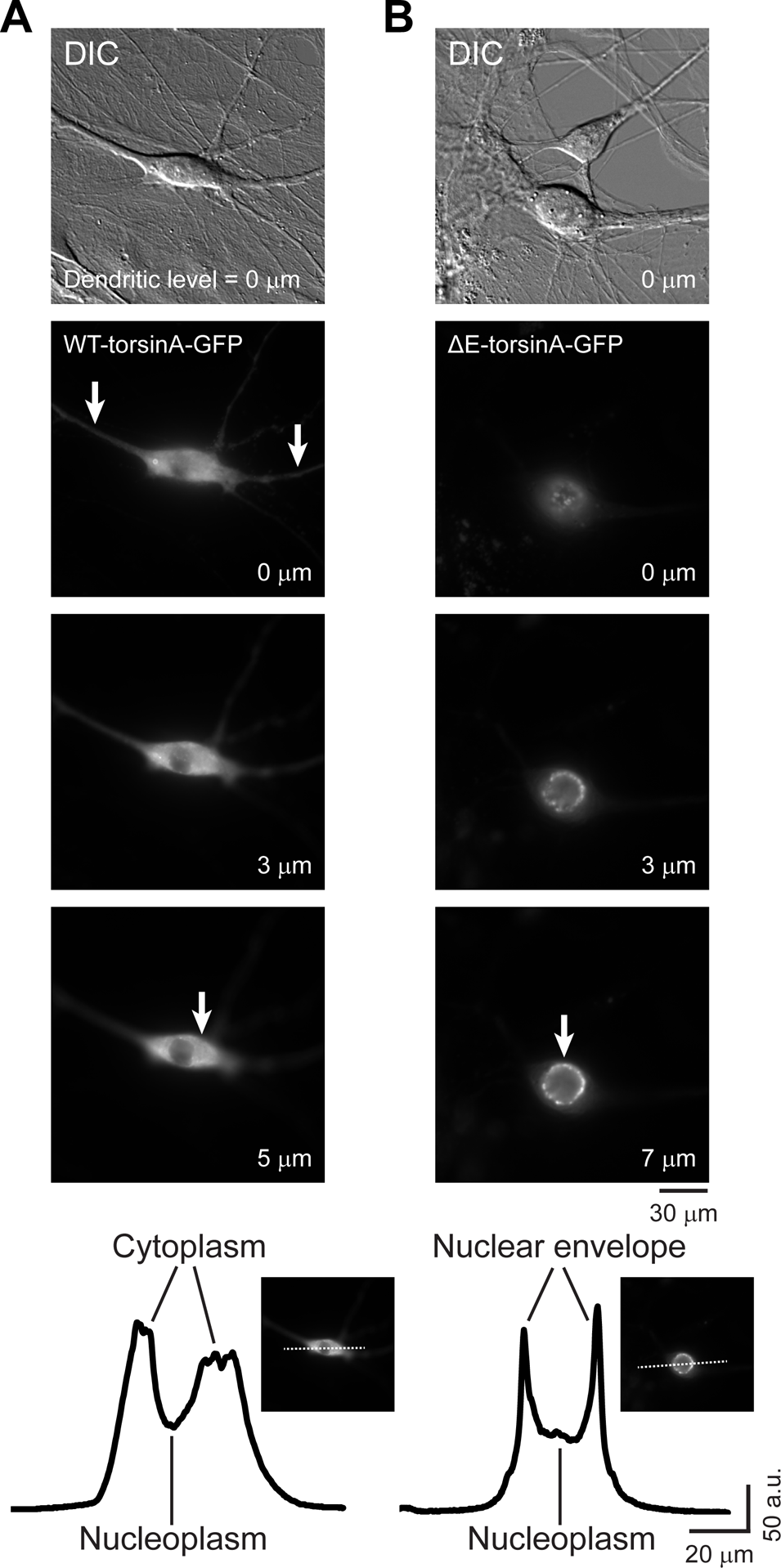
Localization of overexpressed torsinA proteins in hippocampal neurons of wild-type (WT) rats. Cultured neurons transfected with a construct encoding either wild-type (WT)-torsinA (**A**) or ΔE-torsinA (**B**), both tagged with GFP (WT-torsinA-GFP or ΔE-torsinA-GFP, respectively), using a standard calcium-phosphate method. They were observed using a widefield epifluorescence microscope. Differential interference contrast (DIC) imaging was carried out with the focus set at the height of dendrites (0 µm). Three fluorescence images were taken at this and two additional focal levels (at 0, 3, and 5 or 7 μm). Arrows point to diffuse WT-torsinA-GFP fluorescence in the dendrites and soma (**A**), and the GFP signal surrounding the nucleus (perinuclear pattern) (**B**). Line-intensity plots (intensity plotted against pixels along a line) show that the main signal of WT-torsinA-GFP is in the cytoplasm and is diffuse, with weaker signal in the nucleoplasm. The main signal of ΔE-torsinA-GFP is perinuclear, with weaker signal in the nucleoplasm.

We next modified the transfection method to improve efficiency (Jiang and Chen, 2006). In these experiments, signal from the overexpressed WT-torsinA-GFP in the rat hippocampal neurons was diffuse in the cytoplasm (arrows) with additional perinuclear signal (top row, Fig. 2A). In the figure, a single confocal section is used to demonstrate the distribution at the somatic level, and a maximum intensity projection (MIP) is used to demonstrate the distribution throughout the cell, incorporating both the dendritic and somatic signals. More dramatic differences were observed in the case of overexpressed ΔE-torsinA-GFP. It was localized mainly in cytoplasmic inclusion bodies and in the perinuclear region (top row, Fig. 2B). The MIP images reveal a lack of diffuse dendritic and somatic staining. These findings are consistent with previous reports that at high levels of expression, WT-torsinA protein is localized more toward the nuclear envelope, and ΔE-torsinA is localized more to inclusion bodies (Hewett et al., 2000; Kustedjo et al., 2000; Bragg et al., 2004a; Goodchild and Dauer, 2004; Harata, 2014).

**Fig. 2.**
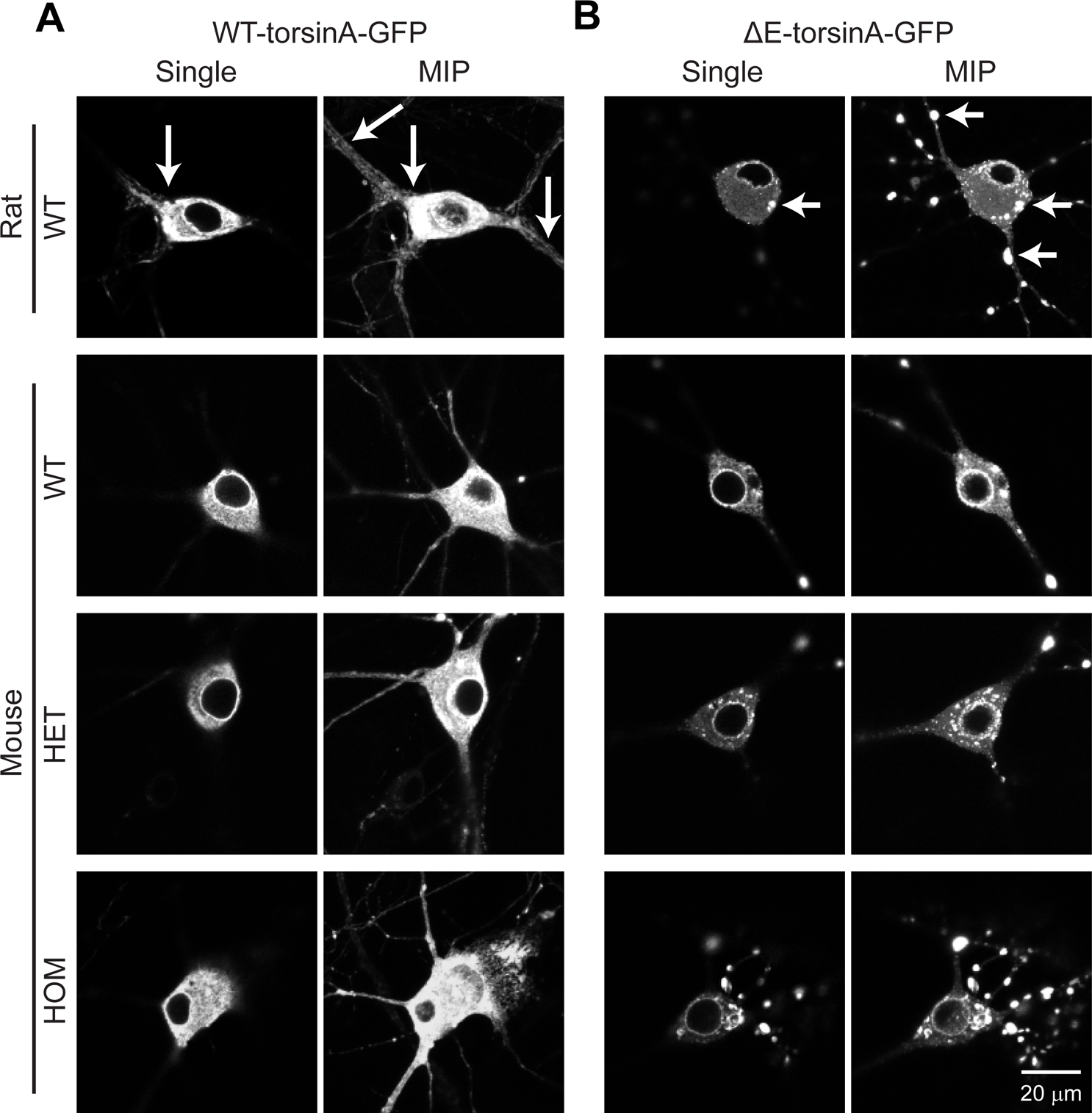
Localization of overexpressed torsinA proteins in hippocampal neurons of rats and ΔE-torsinA knock-in mice. Cultured neurons were transfected with a construct encoding either WT-torsinA-GFP (**A**) or ΔE-torsinA-GFP (**B**). A modification of the calcium-phosphate method in Fig. 1 was used to enhance the efficiency of transfection (Jiang and Chen, 2006). A confocal laser-scanning microscope was used to observe the neurons. Neurons are shown both as single optical sections at the somatic level (Single) and as maximum intensity projection (MIP) images that represent the overall structure at all heights. Neurons were cultured from WT rats (top row), WT mice (2^nd^ row), heterozygous (HET, *Tor1a*^+/ΔE^, 3^rd^ row) or homozygous (HOM, *Tor1a*^ΔE/ΔE^, bottom row) ΔE-torsinA knock-in mice. Arrows point to representative examples of the diffuse distribution of WT-torsinA-GFP in the cytoplasm (**A**), and ΔE-torsinA-GFP in cytoplasmic inclusion bodies (**B**).

The literature also includes reports of interactions between WT- and ΔE-torsinA proteins. The overexpression of both WT- and ΔE-torsinA in the same WT cells has been shown to drive the overexpressed WT form to co-localize with ΔE form in BHK21 cells (Goodchild and Dauer, 2004), HEK293 cells (Torres et al., 2004) and PC6-3 cells (Gonzalez-Alegre et al., 2005). We examined whether such interactions are detectable when one protein is expressed at endogenous levels. The localization of overexpressed proteins in mouse hippocampal neurons was not affected by the endogenous forms, irrespective of whether the host neurons were obtained from WT (*Tor1a*^+/+^), heterozygous (HET, *Tor1a*^+/ΔE^) or homozygous (HOM, *Tor1a*^ΔE/ΔE^) ΔE-torsinA knock-in mice (3 rows at bottom, Fig. 2). More specifically, the location of overexpressed WT torsinA in heterozygous or homozygous neurons was not affected by the presence of ΔE-torsinA protein at the endogenous level. Conversely, the location of overexpressed ΔE-torsinA in WT or heterozygous neurons was not affected by the presence of WT-torsinA protein at the endogenous level. Thus the localization of an overexpressed protein was dictated by the form of the overexpressed protein itself, rather than that of the endogenous proteins in the host cells. In summary, the localization of overexpressed torsinA proteins can differ from that of endogenous proteins.

### Subcellular localization of endogenous torsinA proteins in WT mouse neurons

Immunocytochemistry was used to evaluate the distribution of endogenous torsinA proteins in cultured neurons. The specificity of the primary, polyclonal anti-torsinA antibody used in this study has been tested extensively (see Materials and Methods section). Those tests included the demonstration of a lack of immunoreactivity in the brains of conditional *Tor1a* knock-out mice (Liang et al., 2014). In addition, we showed that immunostaining of cultured rodent hippocampal cells was eliminated when the antibody was preadsorbed with immunizing peptide (Koh et al., 2013).

Given the broad expression of torsinA within the brain (Augood et al., 1999; Shashidharan et al., 2000; Walker et al., 2001), its function is expected to be universally relevant to neurons. First, we tested cultured hippocampal neurons because the hippocampus is among the brain regions that show the highest levels of *Tor1a* transcript (Augood et al., 1998; Augood et al., 1999; Ziefer et al., 2002; Allen_Mouse_Brain_Atlas, 2004) and torsinA protein levels (Walker et al., 2001). Also the predominance of pyramidal neurons with tall somata in hippocampal culture makes it possible to image the neuronal signal with minimal stray signal from the glial cells that lie under the neurons.

Hippocampal neurons of WT mice were stained for endogenous torsinA (Fig. 3A) and images were acquired at three focal levels. Near the surface of the coverslip, glial cells were positive for torsinA (upward arrows in glial layer, labeled 0 µm) and their nuclei were stained with the nuclear Hoechst dye (upward arrowheads). Neuronal signal was out of focus at this height. Two µm higher (away from the coverslip), neuronal torsinA signal was strong and in focus in the soma (downward arrow), and weak in dendrites. This was not at exactly the same height as the neuronal nucleus. Four µm above the glial layer, the neuronal torsinA signal was partly out of focus, but the nucleus was in clear focus (downward arrowhead). Thus, the main signal of the immunoreactive, endogenous torsinA was paranuclear (near the nucleus, but not necessarily surrounding it in a perinuclear ring-like pattern). Minor components were in the nuclear region and dendrites. This pattern differed from the uniform and extensive distribution of overexpressed WT-torsinA (Figs. 1, 2).

**Fig. 3.**
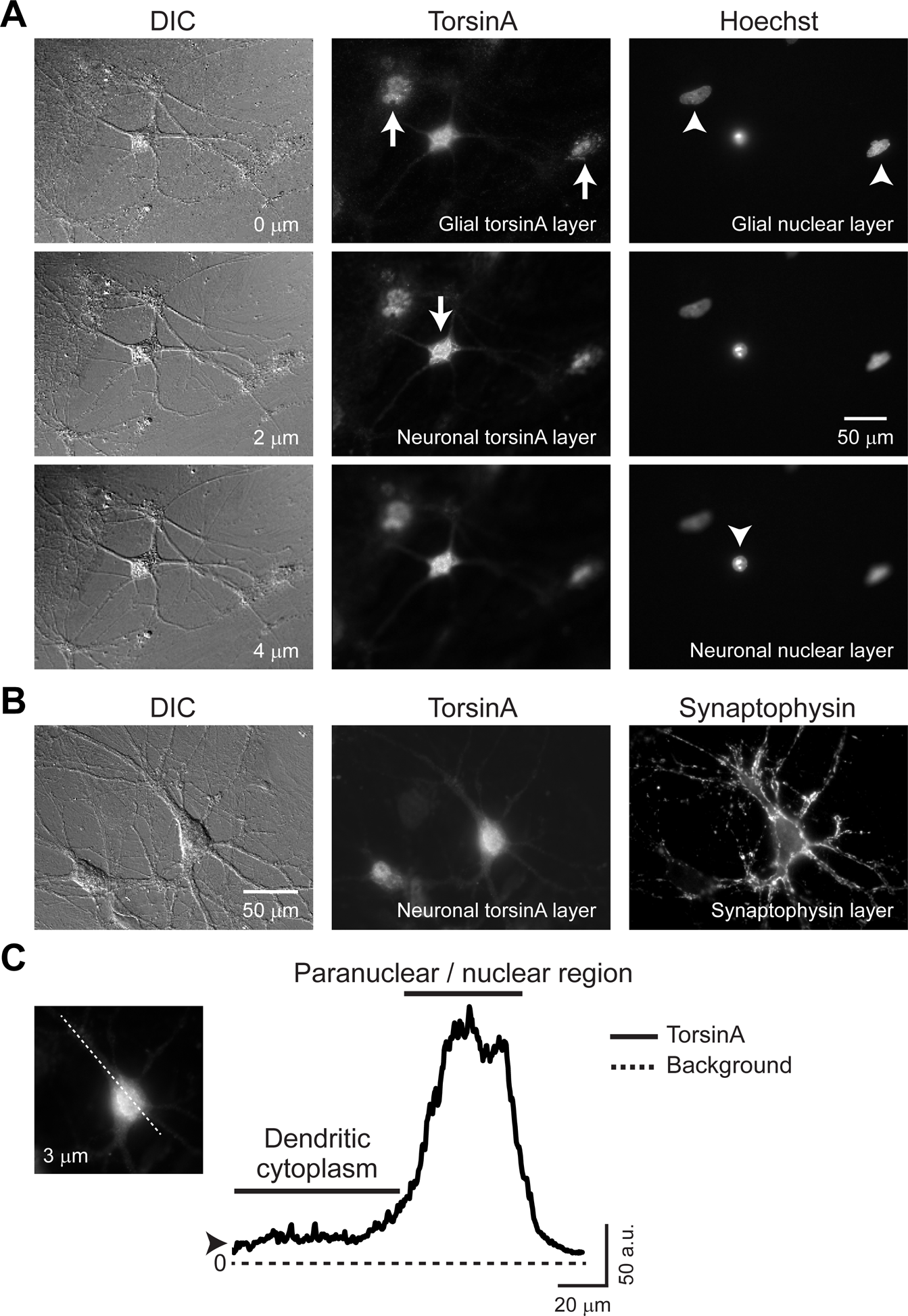
Localization of endogenous torsinA in cultured mouse hippocampal neurons. Hippocampal neurons were cultured from WT mice and stained for torsinA by immunocytochemistry. They were observed using a widefield microscope. **A**) DIC image showing the general morphology of the cells, and fluorescence microscopy showing torsinA signal and nuclei counterstaining with Hoechst 33342 dye. Images were acquired at different focus levels. The glial cell layer is labeled 0 µm. Arrows indicate torsinA signal that is in focus, and arrowheads indicate nuclear signal that is in focus, in neurons (downward) and glial cells (upward). **B**) Neurons co-stained for torsinA and the nerve terminal marker synaptophysin. They were imaged at two different levels of focus. The somatic layer was 3 µm above the dendritic, synaptophysin-positive layer. **C**) Line-intensity plot obtained along the dotted line drawn in the image (inset) shown in panel **B**. Arrowhead indicates the staining intensity in the cytoplasm, and the broken line indicates the background intensity of the image. TorsinA and synaptophysin were imaged using secondary antibodies conjugated with Alexa Fluor 488 and Alexa Fluor 568 dye, respectively.

It is possible that the narrower distribution of endogenous torsinA reflects immaturity of the cells or unhealthy culture conditions. However, in previous studies we have shown that hippocampal neurons cultured under the same conditions express 1) presynaptic proteins: synaptophysin (Koh et al., 2013), vesicular glutamate transporter 1 (VGLUT1) (Kakazu et al., 2012b; Iwabuchi et al., 2013a), glutamic acid decarboxylase 65 kDa (GAD65) (Iwabuchi et al., 2013b), 2) a postsynaptic protein: GluA1 subunit of glutamate receptor of AMPA type (Iwabuchi et al., 2013a), and 3) a dendritic protein: microtubule-associated protein 2 (MAP2) (Kakazu et al., 2012b; Iwabuchi et al., 2014b). Moreover, we showed that they are active with functional recycling of synaptic vesicles and synaptic transmission (Kakazu et al., 2012a; Kakazu et al., 2012b). These are all signs of healthy, well differentiated neurons. We tested the differentiation of WT mouse hippocampal neurons in our cultures, by co-staining for torsinA and synaptophysin (Fig. 3B). At the dendritic layer (0 µm), numerous nerve terminals were present on the dendritic and somatic surfaces and in axons (right panel). TorsinA signal was slightly out of focus (not shown). The main torsinA signal was in clear focus at 3 µm higher (middle panel). The presence of extensive processes studded with synaptophysin-positive nerve terminals indicates that these cultured neurons had already developed presynaptic structures, and that the pattern of the torsinA signal was not due to immaturity of the culture. This result also confirms that immunoreactive torsinA is not detectable in the nerve terminals of cultured neurons, consistent with a previous report (Koh et al., 2013).

Line-intensity analysis at the somatic height shows that the main torsinA signal was in the paranuclear and nuclear region (Fig. 3C). Weak cytoplasmic staining was also present (arrowhead). This pattern clearly differs from that of overexpressed WT-torsinA (Figs. 1, 2).

### Subcellular localization of endogenous torsinA proteins in knock-in mice

Our spatial analysis of torsinA expression was extended to the mutant form using the ΔE-torsinA knock-in mouse model (Goodchild et al., 2005) to address whether the ΔE mutation affects the subcellular localization of this protein. Immuno-staining of WT- and ΔE-torsinA proteins is possible because this antibody detects both forms (Koh et al., 2013).

The cerebral cortex and the striatum were tested in addition to the hippocampus. The cerebral cortex was tested because it is involved in motor control associated with dystonia (Neychev et al., 2011), and conditional *Tor1a* knock-out in this region induces motor abnormalities in mice (Yokoi et al., 2008). The striatum was tested in part because it is involved in motor control and is one of the brain regions whose dysfunction is thought to contribute to dystonia (Pisani et al., 2007; Tanabe et al., 2009; Neychev et al., 2011), and the striatum-specific conditional *Tor1a* knock-out mouse shows motor deficits (Yokoi et al., 2011). The striatum is also one of the brain regions in which subtle structural abnormalities have been found in the heterozygous ΔE-torsinA knock-in mice (Song et al., 2013). Moreover, abnormalities in synaptic transmission involving neurotransmitters glutamate (Martella et al., 2009; Sciamanna et al., 2011; Dang et al., 2012; Grundmann et al., 2012; Sciamanna et al., 2012; Martella et al., 2014) and GABA (Sciamanna et al., 2009) have been reported within the striatum of knock-in mice, transgenic mice and transgenic rats.

The WT-torsinA protein (in WT and HET mice) and ΔE-torsinA protein (in HET and HOM mice) were found to localize in the same manner in neurons of the hippocampus (Fig. 4A), cerebral cortex (Fig. 4B) and striatum (Fig. 4C). TorsinA signal was strong in the somatic paranuclear and nuclear region, and also present in proximal dendrites of some neurons. The torsinA signal in the soma can take on a clustered appearance, differing from the diffuse and uniform distribution expected for ER localization. No differences were noted among neurons of different genotypes. For example, staining for ΔE-torsinA was not perinuclear (ring-like) or typical of cytoplasmic inclusion bodies, the patterns that were common with overexpression (Figs. 1, 2). Co-staining for synaptophysin demonstrated abundant presynaptic nerve terminals, indicating that the synapses in our *in vitro* system were mature.

**Fig. 4.**
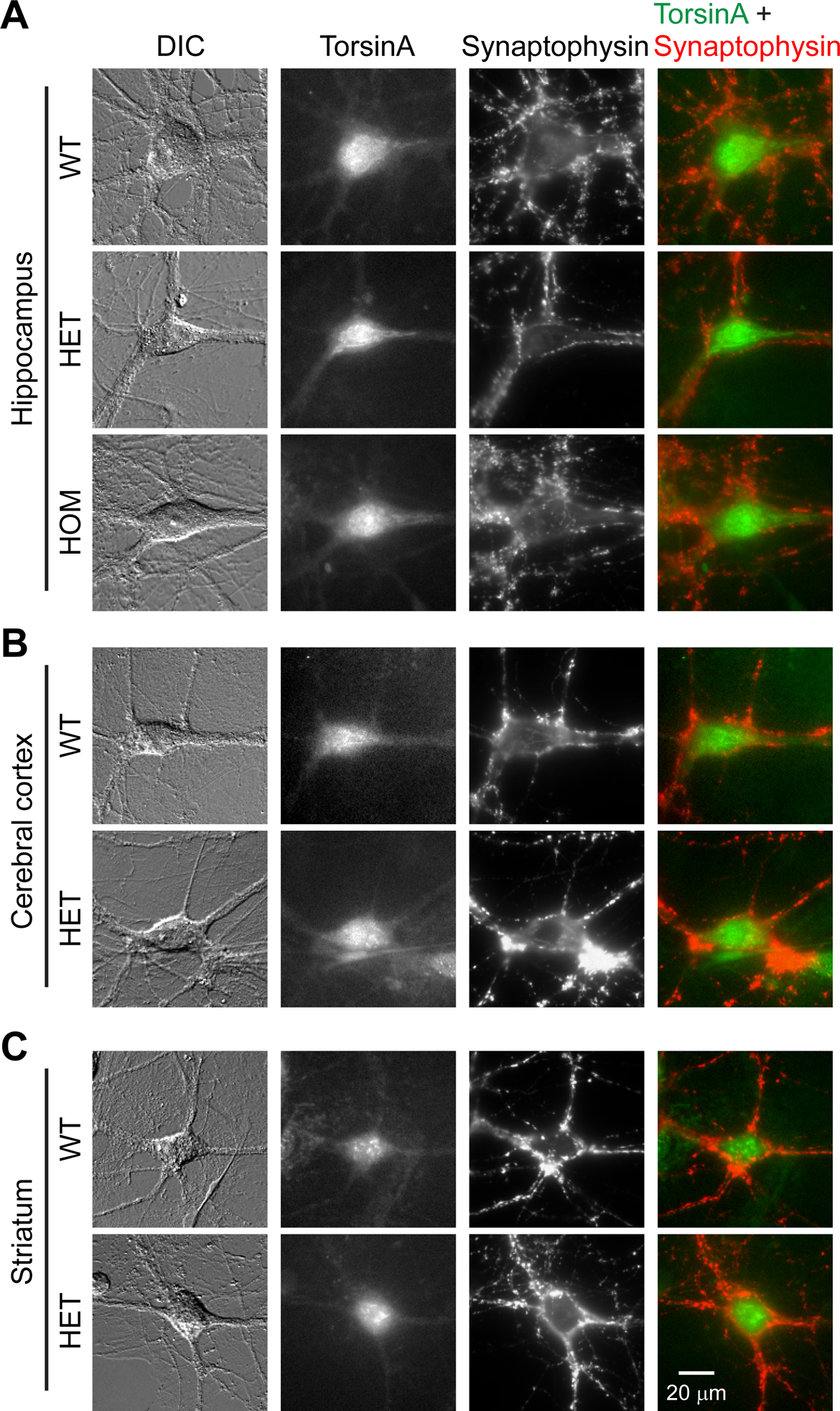
Localization of endogenous torsinA in cultured mouse neurons of different genotypes in three brain regions. Neurons from WT (WT, *Tor1a*^+/+^), heterozygous (HET, *Tor1a*^+/ΔE^) or homozygous (HOM, *Tor1a*^ΔE/ΔE^) ΔE-torsinA knock-in mice were cultured, co-stained for torsinA and synaptophysin by immunocytochemistry, and observed with a widefield microscope. Neurons were obtained from the hippocampus (**A**), cerebral cortex (**B**), and striatum (**C**). DIC and synaptophysin images were acquired at the dendritic focus level, and torsinA images were acquired 2 µm higher. TorsinA and synaptophysin were visualized using secondary antibodies conjugated with Alexa Fluor 488 and Alexa Fluor 568 dye, respectively. There was no detectable difference in the distributions of WT-torsinA (in WT and HET mice) and ΔE-torsinA (in HET and HOM mice).

Thus the distribution patterns of endogenous torsinA were identical in neurons of different genotypes and of different brain regions. Also they were very different from the patterns expected from mis-localized ΔE-torsinA.

### Endogenous torsinA near the Golgi apparatus of WT mouse hippocampal neurons

Strong staining near nucleus indicates that endogenous torsinA is distributed in an organelle other than the ER. In order to test this hypothesis, we tested the torsinA signal for co-localization with the markers of various organelles by immunocytochemistry.

The cultured hippocampal neurons of WT mice were co-stained for torsinA and the *cis*-Golgi marker GM130 (Horton et al., 2005), and imaged by confocal microscopy to provide high spatial resolution (Fig. 5A). The paranuclear torsinA signal co-localized extensively with GM130 signal, with both forming irregular shapes. Line-intensity analysis in the two channels shows that the most intense component of the torsinA signals showed variation similar to that of GM130 signal (asterisks), strongly supporting their co-localization. Within the cytoplasm, the torsinA signal was weak and diffuse (arrowhead).

**Fig. 5.**
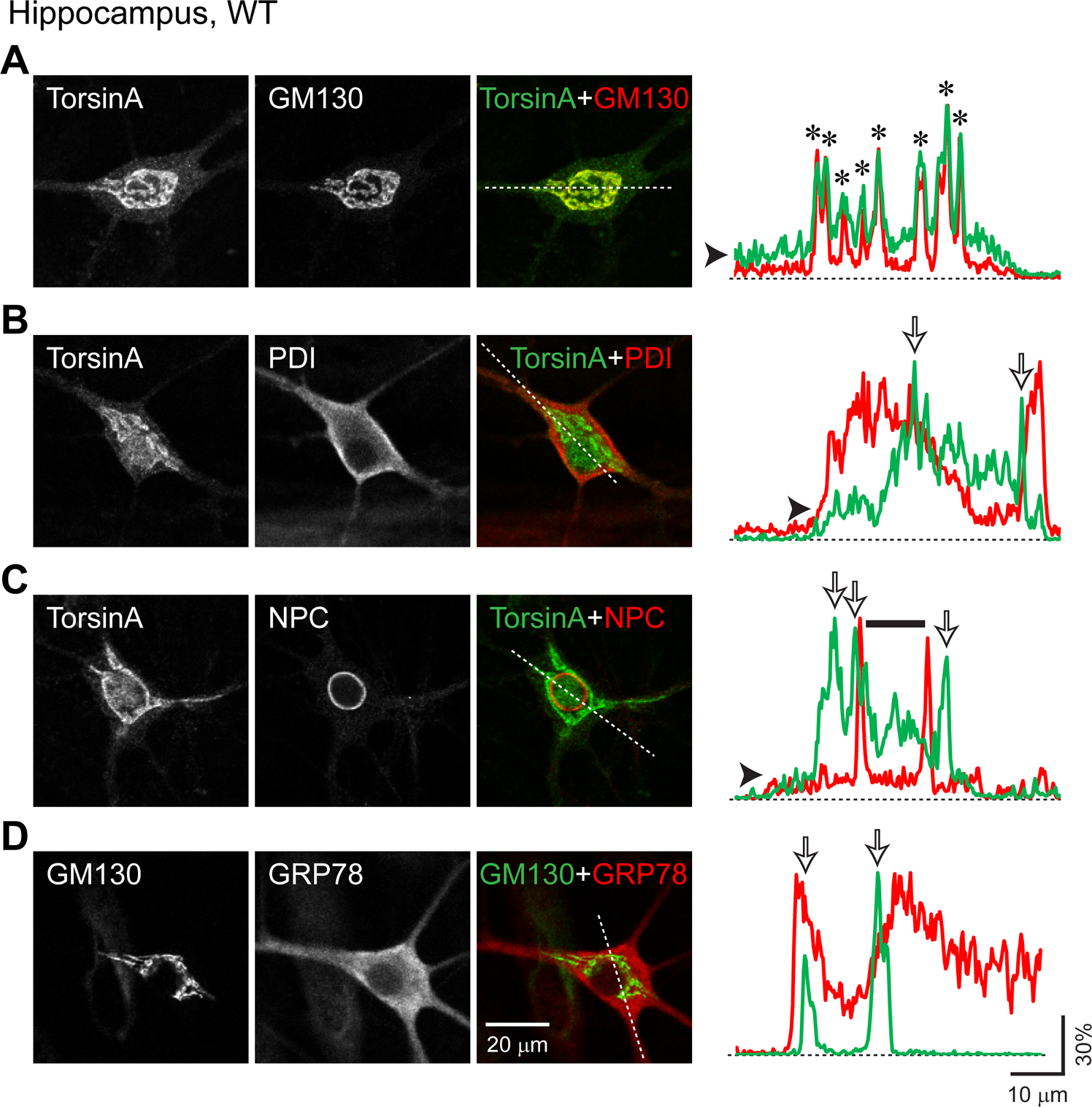
Localization of endogenous torsinA relative to the Golgi apparatus, endoplasmic reticulum (ER) and nuclear envelope, in cultured neurons from the WT mouse hippocampus. WT mouse hippocampal neurons were imaged with confocal microscopy. **A**) A neuron double-stained for torsinA and the *cis*-Golgi marker, Golgi matrix protein of 130 kDa (GM130). **B**) A neuron double-stained for torsinA and the ER marker, protein disulfide isomerase (PDI). **C**) A neuron double-stained for torsinA and the nuclear envelope marker, nuclear pore complex (NPC) protein. **D**) A neuron double-stained for GM130 and the ER marker, glucose-regulated protein of 78 kDa / binding immunoglobulin protein (GRP78/BiP). TorsinA and GRP78 were visualized using Alexa Fluor 488 (green channel), and GM130, PDI and NPC using Alexa Fluor 568 (red channel). In panel **D**, the colors in the pseudo-color images were swapped. In each grayscale image, the intensity scale was adjusted so that the minimum and maximum correspond to the 0 and 255 values on an 8-bit scale. A dotted line was drawn in the overlaid image of the two channels, and was used to generate intensity plot for each channel. The traces were normalized, with the peak intensity as 1 and the background level as 0. They were overlaid and color-coded as the antigens in the overlaid image. Asterisks indicate extensive colocalization of torsinA with GM130. Open arrows indicate torsinA peaks that partially colocalized with PDI and NPC, and GM130 peaks that partially colocalized with PDI. Arrowheads indicate diffuse torsinA staining in the cytoplasm. The dotted lines indicate the background intensity. Horizontal bar in **C** indicates nucleoplasmic torsinA staining, defined as staining interior to the ring of NPC signal. The main signal of endogenous WT-torsinA was in the Golgi apparatus.

TorsinA did not co-localize extensively with the ER marker PDI (Fig. 5B). Although the main torsinA signal overlapped partially with that of PDI (open arrows), in many other regions one signal was weak and the other strong. Also, the torsinA signal was clustered in small regions whereas the PDI signal was diffuse throughout the cell. However, the weak and diffuse component of the torsinA signal in the cytoplasm seemed to co-localize with that of PDI (arrowhead). This localization pattern differs from that of overexpressed WT-torsinA, which co-localized extensively with the ER markers PDI (O’Farrell et al., 2002; Bragg et al., 2004b; Naismith et al., 2004; Misbahuddin et al., 2005; Jungwirth et al., 2010; Nery et al., 2011; Hettich et al., 2014), GRP78/BiP (Kustedjo et al., 2000) and KDEL (Goodchild and Dauer, 2004; Calakos et al., 2010; Maric et al., 2011).

We also evaluated the distribution of endogenous torsinA within the nuclear envelope because the overexpressed torsinA proteins, both the WT and ΔE forms, colocalize with the nuclear envelope marker nucleoporin (Bragg et al., 2004a). However, the main torsinA signal differed from the ring-like staining of nucleoporins detected by the anti-NPC antibody (Fig. 5C). A similar relationship was observed with antibody for another nuclear envelope marker lamin (data not shown). In some neurons, additional torsinA staining was present in the nucleoplasm (horizontal bar in line-intensity analysis).

### Spatial relationship between the Golgi apparatus and ER

If the most intense component of the torsinA signal overlaps partially with that for the ER (Fig. 5B), is this contradictory to torsinA being preferentially localized near Golgi apparatus (Fig. 5A)? In order to answer this question, the hippocampal neurons of WT mice were co-stained for markers of the Golgi (GM130) and ER (GRP78/BiP) (Fig. 5D). The signals colocalized partially near the nucleus. However, the spatial extent of GM130 staining was more limited than that of GRP78, as reported for hippocampal neurons stained with other markers of Golgi and ER (Krijnse-Locker et al., 1995; Horton and Ehlers, 2003). Overall, GM130 was colocalized with GRP78, but the converse is not true. The Golgi apparatus is surrounded by the ER, with the minimum distance separating them being tens of nanometers (Peters et al., 1991), whereas the optical resolution for discerning two separate points is limited to 200-300 nm (Rayleigh criterion) (Egner and Hell, 2006; Inoue, 2006). Thus the seemingly overlapping signals in the paranuclear region are due to the limitations of optical resolution, and the objects are distinct (Golgi and ER). Namely, the staining patterns for the ER and Golgi apparatus will not necessarily be non-overlapping in the paranuclear region. Therefore the results in Figs. 5A and 5B are not contradictory.

### Localization of the main torsinA signal near Golgi apparatus in cultured hippocampal neurons at multiple heights

The images in Fig. 5 were acquired at the heights at which the neuronal torsinA signal was strongest. Fig. 6 shows that torsinA and GM130 signals colocalized at different levels of a z-stack. This result is consistent with the main torsinA signal being present near the *cis*-Golgi.

**Fig. 6.**
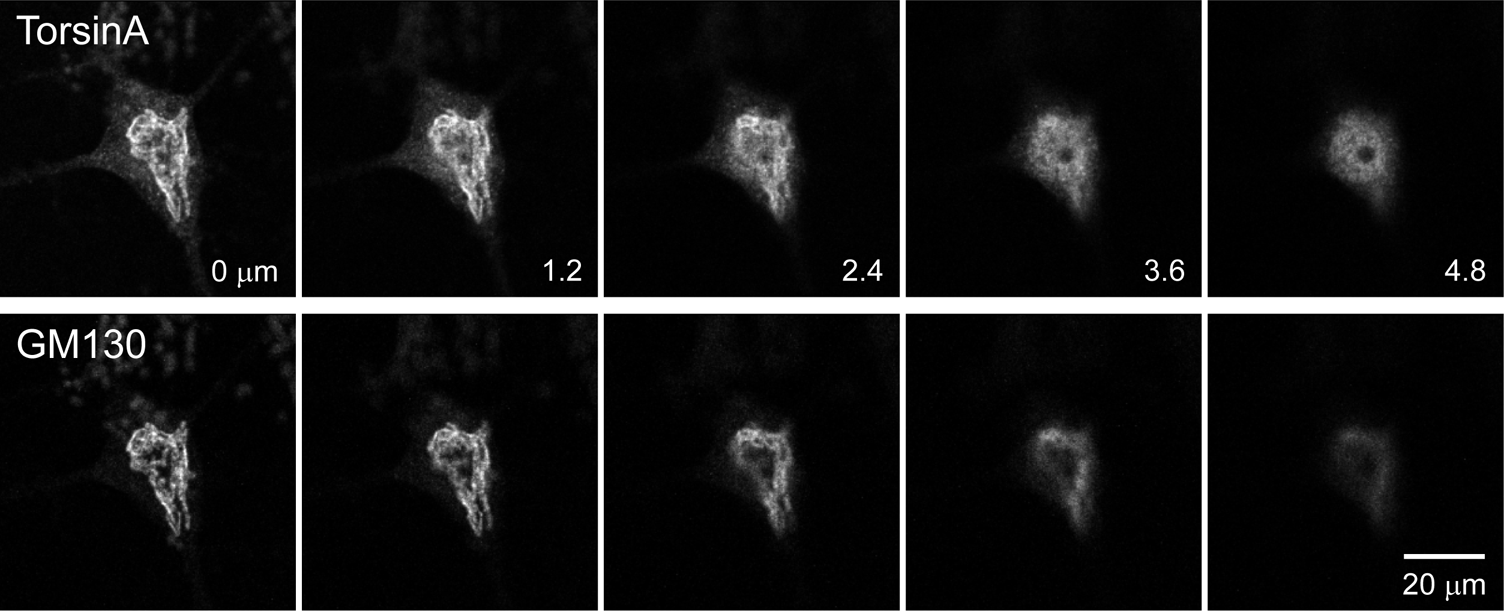
Co-localization of endogenous torsinA and Golgi signals in cultured hippocampal neurons of WT mice. TorsinA and GM130 images are shown at different levels of a confocal z-stack. The numbers represent the height in μm from the dendritic layer. The torsinA and GM130 signals colocalized at different heights. A weaker signal was also present in the nucleoplasm.

### Localization of endogenous torsinA near the Golgi apparatus in cultured hippocampal neurons of mutant mice

We next assessed which organelles the torsinA proteins in mutant mice localize to, using cultured hippocampal neurons. In both heterozygous (Fig. 7, top half) and homozygous neurons (Fig. 7, bottom half), the main torsinA signal colocalized with the GM130 signal, and only weakly with the ER marker PDI or the nuclear envelope marker NPC. The torsinA signal did not take on a perinuclear ring-like or cytoplasmic inclusion-like pattern as observed when the ΔE-torsinA protein was overexpressed in the same culture system (Fig. 2B). Essentially, the distribution of torsinA signal in the mutant neurons was identical to that of in WT neurons (Fig. 5). Thus endogenous torsinA, whether in the WT or ΔE form, is present predominantly near the Golgi apparatus of hippocampal neurons.

**Fig. 7.**
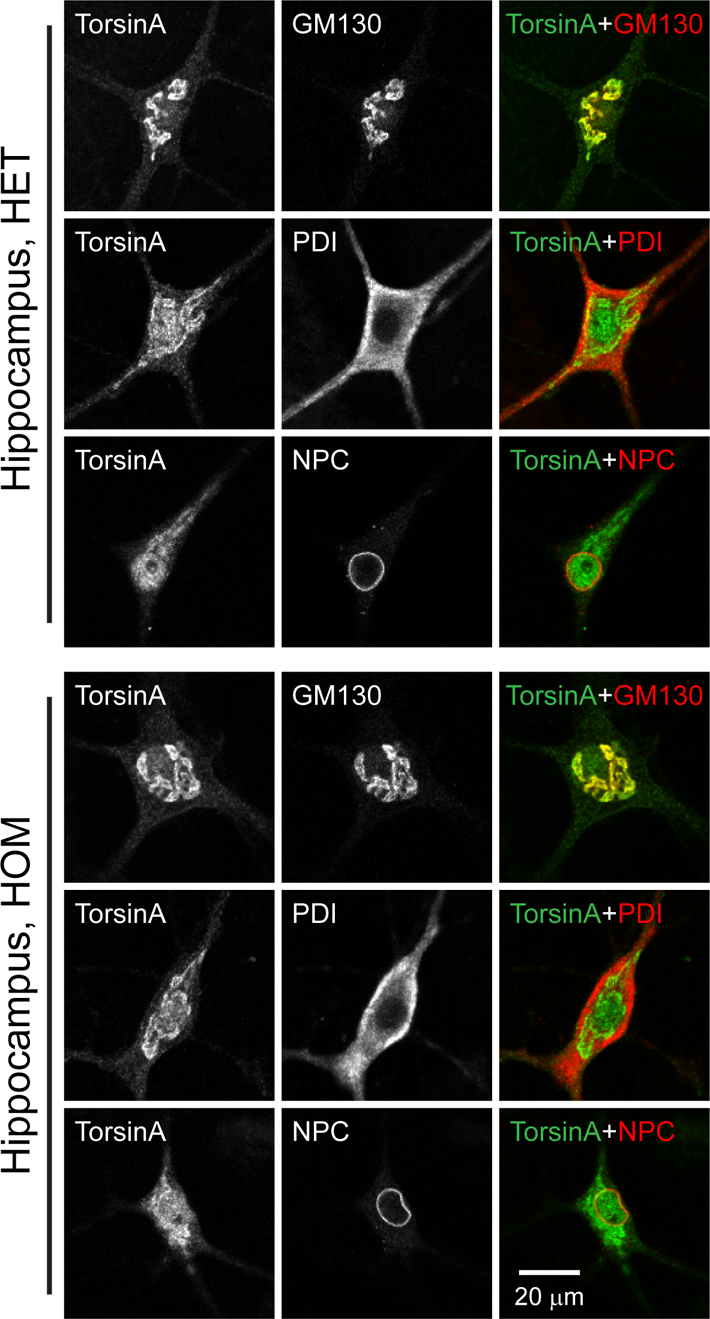
Localization of endogenous torsinA in cultured hippocampal neurons of heterozygous and homozygous mutant mice. Hippocampal neurons were imaged by confocal microscopy. **Top**) Neurons of HET mice, double-stained for endogenous torsinA, and GM130, PDI or NPC. **Bottom**) Neurons of HOM mice, double-stained as in panel A. The distributions of torsinA proteins were the same as in WT mice (Fig. 5).

### Localization of torsinA near the Golgi apparatus in cultured cerebral cortical and striatal neurons

The immunocytochemical analysis described above for hippocampal neurons was applied to cultured cerebral cortical (Fig. 8) and striatal (Fig. 9) neurons of the three genotypes. As in the case of the hippocampal neurons, differences in torsinA protein distribution were not detected among the WT, heterozygous and homozygous ΔE-torsinA knock-in mice. The main signal was present near the Golgi apparatus, accompanied by weak nuclear envelope and ER staining. Mutant neurons did not show perinuclear ring-like staining or cytoplasmic inclusions.

**Fig. 8.**
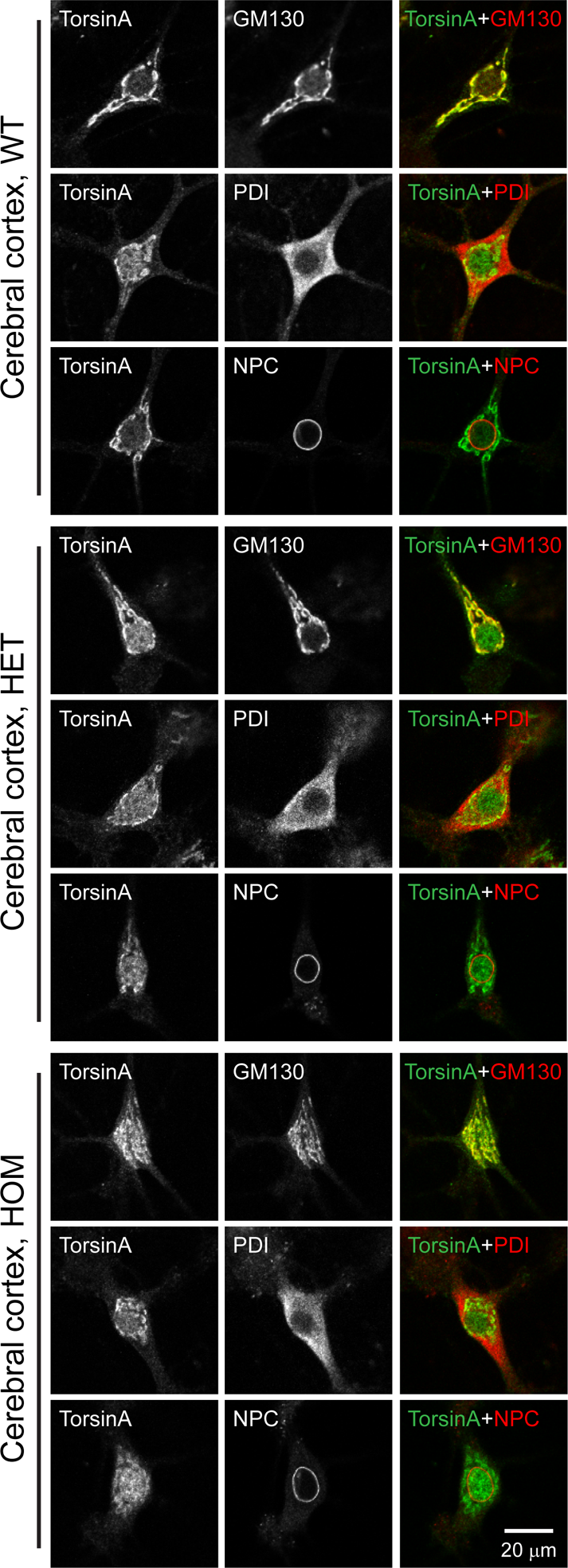
Localization of endogenous torsinA in cultured cerebral cortical neurons of WT, heterozygous and homozygous-mutant mice. **Top**) Neurons of WT mice, double-stained for endogenous torsinA, and GM130, PDI or NPC. **Middle**) Neurons of the HET mice, double-stained the same way as in panel A. **Bottom**) Neurons of the HOM mice, double-stained as in panels A, B. Neurons were imaged by confocal microscopy. The torsinA distributions were the same in the mice of all three genotypes.

**Fig. 9.**
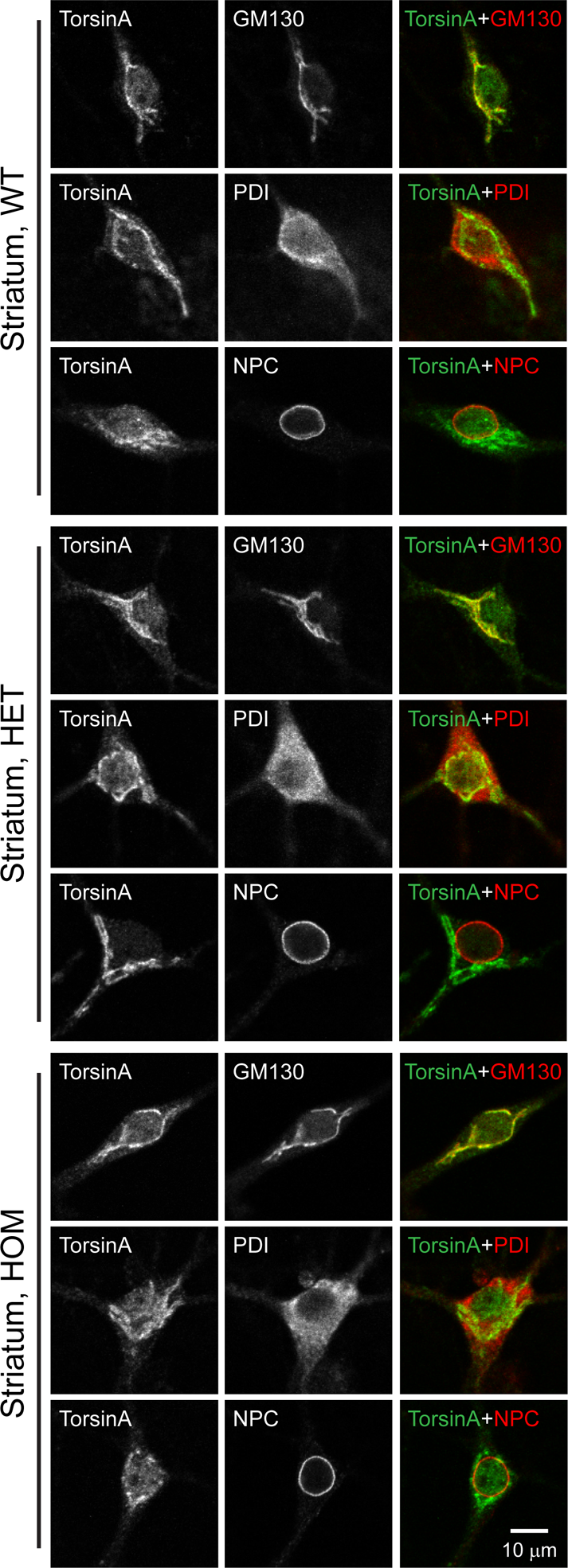
Localization of endogenous torsinA in cultured striatal neurons of WT, heterozygous and homozygous-mutant mice. **Top**) Neurons of WT mice, double-stained for endogenous torsinA, and GM130, PDI or NPC. **Middle**) Neurons of the HET mice, double-stained the same way as in panel A. **Bottom**) Neurons of the HOM mice, double-stained as in panels A, B. Neurons were imaged by confocal microscopy. The torsinA distributions were the same in the mice of all three genotypes.

### TorsinA localization near the Golgi apparatus in cultured rat hippocampal neurons

To exclude the possibility that mouse neurons are affected inappropriately during culture, rat neurons were also tested. The hippocampal neurons of WT rats were co-stained for endogenous torsinA and compartment markers (Fig. 10). The results with GM130, PDI and NPC were identical to those for the mouse hippocampal, cerebral cortical and striatal neurons (Figs. 5, 7-9). The relationship between GM130 and GRP78 (Fig. 10, bottom) was also the same as in WT mouse hippocampal neurons (Fig. 5D). Thus the main torsinA signal was localized near the Golgi apparatus in neurons of two rodent species.

**Fig. 10.**
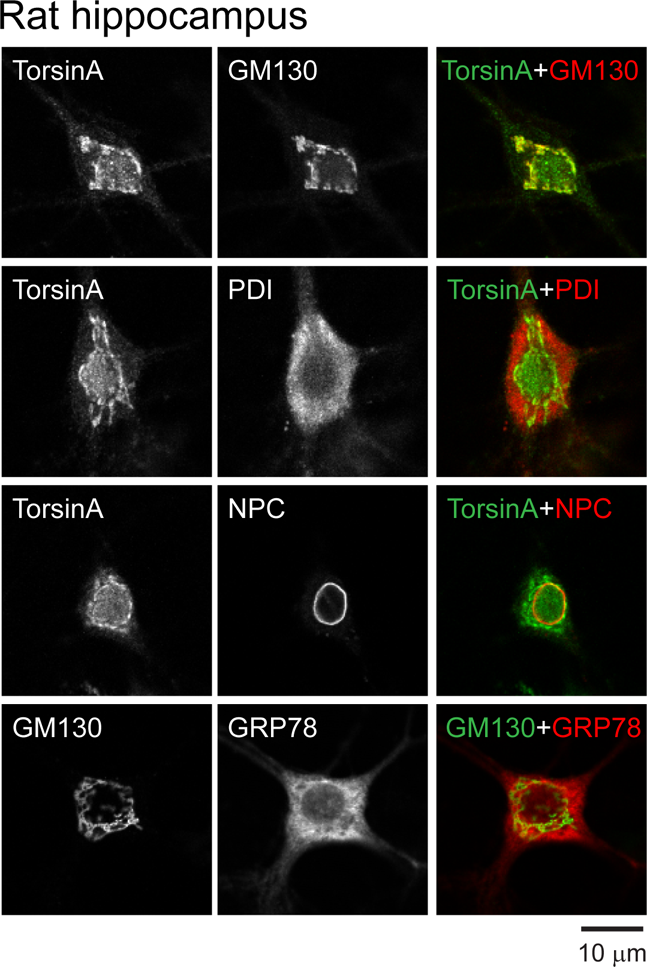
Localization of endogenous WT-torsinA in cultured neurons from the WT rat hippocampus. Analysis was the same as for WT mouse hippocampal neurons (Fig. 5). Neurons were double-stained for endogenous torsinA, and GM130 (1^st^ row), PDI (2^nd^ row) or NPC (3^rd^ row). Neurons were also double-stained for GRP78 and GM130 (4^th^ row). They were imaged by confocal microscopy. The main signal of endogenous WT-torsinA in rat neurons was localized to the Golgi apparatus, as in mouse neurons.

### Relative intensity of endogenous torsinA staining in subcellular compartments

The intensity of endogenous torsinA staining in the Golgi apparatus, ER, nuclear envelope and nucleoplasm was quantified (Fig. 11A), with the compartments identified by the markers used in the above-described experiments. ROI’s were assigned in the channels of individual compartments and transferred to the torsinA channel, and the staining intensity therein was measured. Background intensity was subtracted from the torsinA image beforehand. This non-thresholded analysis of torsinA staining provides unbiased measurement of signal intensity. The intensity of torsinA signal in the subcellular compartments was normalized to general staining in the cytoplasm (ER), because WT-torsinA is thought to be present predominantly in the ER in overexpression studies, and also because ER staining can be used to correct for cell thickness.

**Fig. 11.**
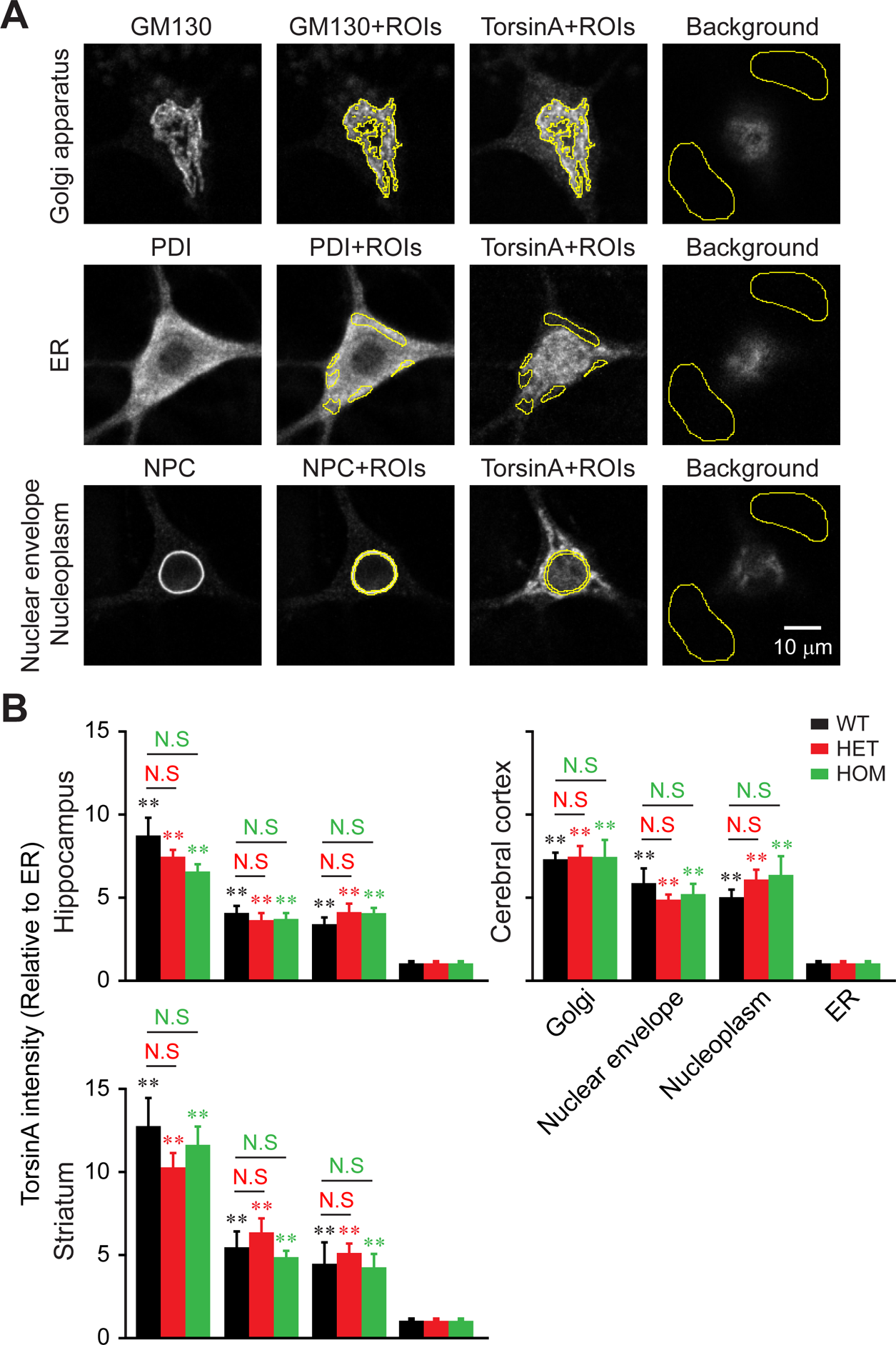
Levels of endogenous torsinA in various subcellular compartments. **A**) Method for measuring torsinA intensity in the Golgi, ER, nuclear envelope and nucleoplasm. Cells were co-stained for torsinA and either GM130 (top), PDI (middle) or NPC (bottom). The regions of interest (ROI’s) for the Golgi apparatus were assigned in the GM130 channel. The ROI’s for the ER were assigned in the PDI channel. The ROI’s for the nuclear envelope and nucleoplasm were assigned based on the NPC-positive ring, as the area within the NPC-positive ring and the area interior to the ring, respectively. These ROI’s were transferred to the torsinA channel. The torsinA pixel intensities in each compartment were then averaged, providing a single value for each cell. These values were used for numerical analysis, after subtracting the averaged background intensity. The background signal was measured from the dimmest regions within the highest image in a z-stack (i.e. a non-cellular region in an image furthest from coverslip level). The ER measurement excluded the Golgi apparatus. The nucleoplasmic measurement excluded nucleoli. **B**) Quantitation of endogenous torsinA in neurons of the hippocampus, cerebral cortex and striatum, obtained from WT, HET and HOM mice. In each panel, the y axis represents pixel intensity normalized to the average ER intensity in each neuron. Imaging conditions were the same for all samples within a single brain region, allowing statistical comparisons of staining intensity. The conditions differed by brain region. TorsinA intensities differed significantly in other compartments vs. the ER (**, p<0.001, n=10-18 neurons / brain region). There were no statistically significant genotypic differences in torsinA intensity between HET or HOM vs. WT for any compartment or any brain region (N.S., p>0.05, n=10-18 neurons in all comparisons).

The relative intensity in the Golgi apparatus was high, ranging between 6.6-12.7 in the 3 brain regions (Fig. 11B). Less prominent expression was observed at the nuclear envelope (based on colocalization with NPC) and the nucleoplasm (interior to the nuclear envelope), with the relative intensities ranging between 3.7-6.5, and 3.4-6.4, respectively. These differences from the ER were statistically significant (**, p<0.001, n=10-18 cells / brain region). There was no genotype-associated difference in the expression levels in any of the 3 brain regions examined (not significant, NS, HET or HOM vs. WT values, p>0.05, n=10-18 cells / brain region).

### Heterogeneity in torsinA staining patterns

Previous reports had indicated that endogenous torsinA is located in the somata of WT rat brain neurons *in situ*, and additionally in the proximal dendrites of some, but not all, neurons (Walker et al., 2001). These neurons included hippocampal neurons. We examined the variability of torsinA staining in the cultured hippocampal neurons, especially in the proximal dendrites. We found that torsinA was present in the somata of all neurons of WT rats and mice of the three genotypes (Fig. 12). However, it was present in the proximal dendrites of only a limited number of neurons (arrows), consistent with the report *in situ* (Walker et al., 2001). This extension of the main torsinA signal into dendrites is consistent with the presence of Golgi apparatus in the dendrites (dendritic Golgi outposts) in some rodent hippocampal and cerebral cortical neurons *in situ* and *in vitro* (Horton et al., 2005; Matsuki et al., 2010; Mitchell et al., 2018). Thus variable torsinA staining in proximal dendrites may reflect the presence or absence of dendritic Golgi outposts.

**Fig. 12.**
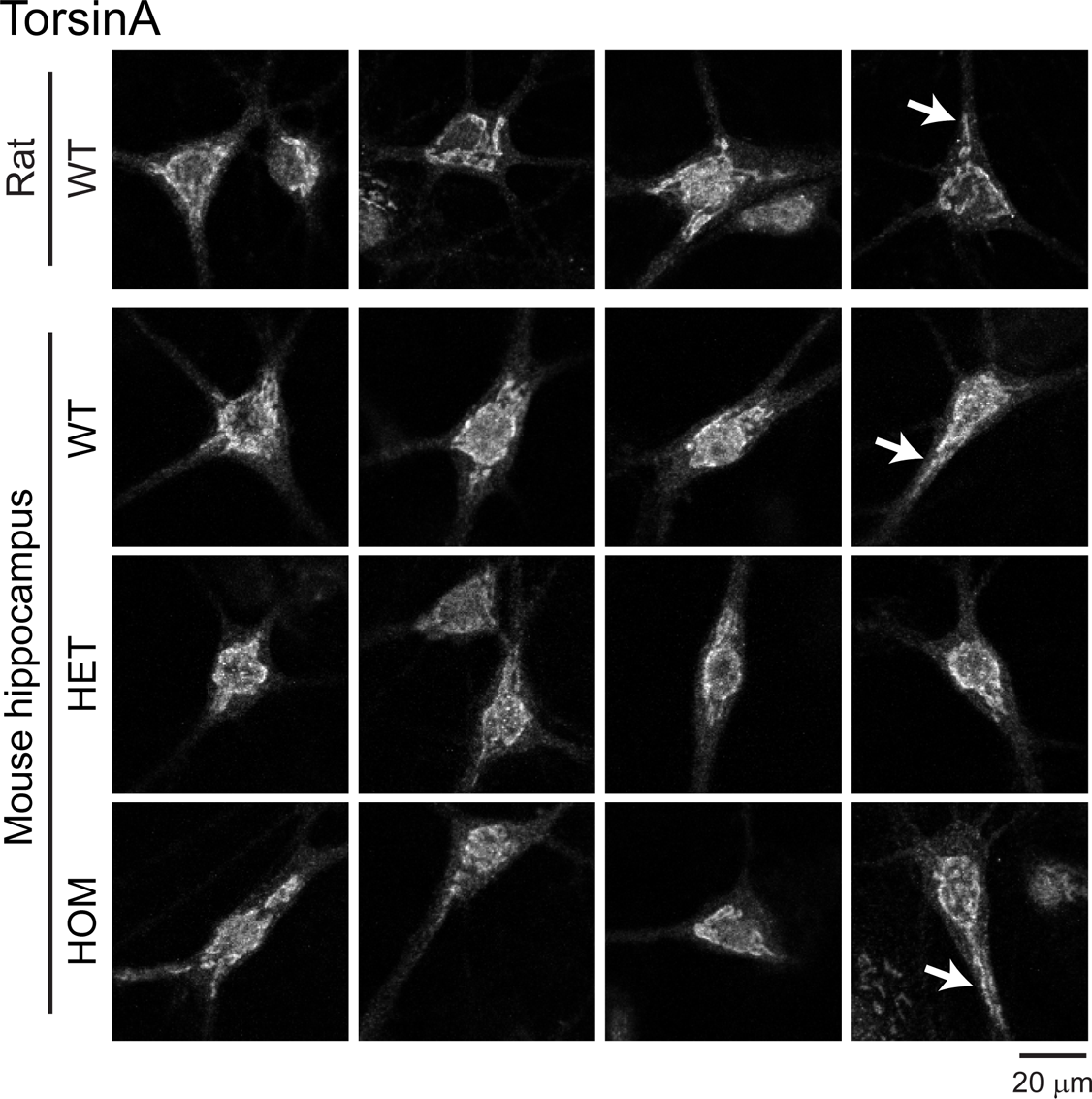
Heterogeneity in torsinA staining in cultured hippocampal neurons. A compilation of torsinA images from WT rat (1^st^ row), WT (2^nd^ row), HET (3^rd^ row) and HOM (4^th^ row) mouse hippocampal neurons. In a small population of neurons, the proximal dendrites were positive (arrows), consistent with the presence of dendritic extensions of the Golgi apparatus.

## DISCUSSION

Here we report the first evidence that the signal of endogenous torsinA protein localizes near the Golgi apparatus in neurons. This was the case in cultured neurons obtained from three brain regions of WT (*Tor1a*^+/+^), HET( *Tor1a*^+/ΔE^) and HOM (*Tor1a*^ΔE/ΔE^) mice, as well as in neurons obtained from WT rat. These results were different from the localizations demonstrated by overexpressed torsinA.

### Distributions of endogenous vs. exogenously overexpressed torsinA proteins

A prevailing model for torsinA localization posits that the WT-torsinA is distributed diffusely throughout the ER, but that ΔE-torsinA is mis-localized to the nuclear envelope and causes abnormalities in its ultrastructure (Harata, 2014). This model is based largely on studies in which torsinA was overexpressed in cultured WT cells of many types, both neurons and non-neuronal cells, and from different species. The non-neuronal cell types include human embryonic kidney epithelial HEK293 cells (Kustedjo et al., 2000), human glioma Gli36 cells (Bragg et al., 2004a), and human osteosarcoma U2OS cells (Kock et al., 2006). Neural cells include human neuroblastoma SH-SY5Y cells (Misbahuddin et al., 2005), mouse neural line CAD cells (Hewett et al., 2000), rat pheochromocytoma PC12 cells (Naismith et al., 2004), PC6-3 cells (Gonzalez-Alegre and Paulson, 2004), and mouse cerebral cortical neurons (Nery et al., 2011). Similar results were reported in central neurons of transgenic mouse (Goodchild and Dauer, 2004) and transgenic rat (Grundmann et al., 2012) that overexpress the WT- and ΔE-torsinA proteins. We confirmed these results by overexpressing the proteins in central neurons (Figs. 1, 2).

Despite the extensive body of literature regarding the above-described hypothesis, there is a general concern that overexpressed proteins can be mis-targeted. For example, endogenous Kv7 potassium channels are localized mainly in a specific compartment of axons (axon initial segment), whereas overexpressed proteins are detected in the ER of somata and dendrites (Rasmussen et al., 2007). Moreover, the previous findings regarding overexpressed torsinA may not accurately reflect the conditions in DYT1 dystonia (*TOR1A*^+/ΔE^), for the following reasons.

First, the level of torsinA expression controls its localization. Thus it is difficult to extrapolate torsinA localization at endogenous levels based on overexpression data. This was demonstrated when WT- or ΔE-torsinA was overexpressed in BHK21 cells (Goodchild and Dauer, 2004) and CHO cells (Naismith et al., 2004), and when the expression level was controlled by a tetracycline-inducible system in Gli36 cells (Bragg et al., 2004a). Specifically, ΔE-torsinA was localized in the ER and nuclear envelope at low expression levels and in intracellular inclusions at high expression levels, whereas WT-torsinA remained in the ER and, to some extent, in the nuclear envelope.

Second, the level of torsinA expression is critical to the nuclear-envelope abnormality. In neurons from mouse brain, this phenotype arises in the absence of two WT alleles, i.e., in *Tor1a*^ΔE/ΔE^ and *Tor1a*^−/−^ mice, but not in the presence of at least one WT allele, i.e., in *Tor1a*^+/+^, *Tor1a*^+/ΔE^ and *Tor1a*^+/−^ mice (Goodchild et al., 2005). The nuclear-envelope abnormality also arises even when WT-torsinA is overexpressed in WT cultured cells (Kim et al., 2010) or in transgenic mice (Grundmann et al., 2007). Thus, extreme up- or down-regulation of WT torsinA can cause abnormalities of the nuclear envelope.

Third, overexpression of torsinA can induce functional changes in neurons. In transgenic mice overexpressing even WT-torsinA, there were changes in dopamine release and animal behaviors relative to non-transgenic controls (Grundmann et al., 2007; Page et al., 2010).

Fourth, the torsinA proteins are not overexpressed in the fibroblasts of DYT1 dystonia patients (Goodchild et al., 2005), or in the striatum (Yokoi et al., 2010), hippocampus (Yokoi et al., 2013) and whole brain (Yokoi et al., 2012) of heterozygous ΔE-torsinA knock-in (*Tor1a*^+/ΔE^) mice.

Fifth, in patients with DYT1 dystonia, torsinA distribution in the brain (Walker et al., 2002a; Rostasy et al., 2003) is similar to that in normal subjects (Shashidharan et al., 2000; Walker et al., 2002a; Rostasy et al., 2003) and WT rats (Shashidharan et al., 2000; Walker et al., 2001). TorsinA is present in the cytoplasm of the soma as well as in some proximal dendrites and the nucleoplasm, does not co-localize extensively with the ER, shows no intracellular inclusions, and has a distribution that appears “to differ from that seen in cell culture transfection studies” (Walker et al., 2002a).

Sixth, there is a discrepancy between the predicted localization and the findings from WT rodent and human brains. Had the WT-torsinA been diffusely and evenly distributed in the ER, the torsinA distribution would have mimicked the distribution of ER proteins, e.g. inositol-1,4,5-trisphosphate receptor and ryanodine receptor. Nevertheless, torsinA localization in somata and proximal dendrites in human and rat brains (Shashidharan et al., 2000; Konakova et al., 2001; Walker et al., 2001; Walker et al., 2002a; Rostasy et al., 2003) is strikingly different from the pronounced neuropil staining observed for ER proteins (Hertle and Yeckel, 2007).

Therefore, it seems necessary to exercise caution when localization of overexpressed torsinA is extrapolated to that of the endogenously expressed protein.. Moreover, the localization of endogenous torsinA has not been addressed extensively (Harata, 2014), although its importance was pointed out in one of the first studies to localize overexpressed torsinA (Kustedjo et al., 2000), and also in later work on the endogenous (Walker et al., 2002a; Xiao et al., 2004) and exogenous distribution (Baptista et al., 2004).

### WT- and ΔE-torsinA proteins, expressed at endogenous levels, are localized primarily near the Golgi apparatus

Localization of overexpressed torsinA has been tested using immunocytochemical markers of the ER (Hewett et al., 2000; Kustedjo et al., 2000; Gonzalez-Alegre and Paulson, 2004; Naismith et al., 2004; Misbahuddin et al., 2005; Nery et al., 2011), nuclear envelope (Bragg et al., 2004a; Gonzalez-Alegre and Paulson, 2004), and lysosomes (Calakos et al., 2010). However, localization to the Golgi apparatus has not been tested except in two reports using overexpression systems. One of these analyses showed that overexpressed ΔE-torsinA did not colocalize with GM130 in differentiated, neuron-like CAD cells (Hewett et al., 2000). The other reported that, when the N-terminal domain (amino acids 26-43 out of 332) was deleted from the mature torsinA and the protein was overexpressed in cultured osteosarcoma U2OS cells, it accumulated in the Golgi apparatus and was secreted to the extracellular solution (Vander Heyden et al., 2011). It is important to note that a proteomics study of WT mouse liver identified endogenous torsinA in the Golgi apparatus (Foster et al., 2006). Unfortunately, this finding was later dismissed as contamination from the ER (Gilchrist et al., 2006), simply based on the notion that overexpressed WT-torsinA had been found exclusively in the ER (Hewett et al., 2003). Interestingly, the possibility that torsinA is associated with the Golgi was discussed in an immunohistochemical study of human testis (Serrano et al., 2019).

In contrast to the data from overexpression systems, those from analyses of endogenous torsinA proteins have been inconsistent. In the studies carried out *in vitro*, WT-torsinA was present in the cell body (Hewett et al., 2003; Gonzalez-Alegre et al., 2005). It was also detected in neurites and growth cones of immature cells (Ferrari-Toninelli et al., 2004; Kamm et al., 2004; Granata et al., 2008; Granata et al., 2011), but not the axonal shafts and nerve terminals of more mature neurons (Koh et al., 2013). In studies of neurons *in vivo*, the results were more variable. The majority of reports found that WT-torsinA is distributed diffusely in the cytoplasm of somata and proximal dendrites. However, the results also range from strong staining of axons and nerve terminals (Konakova et al., 2001; Konakova and Pulst, 2001; Walker et al., 2002b; Rostasy et al., 2003), to neuropil staining in the absence of soma or dendrite staining (Augood et al., 2003) and staining of the nucleoplasm (Shashidharan et al., 2000; Walker et al., 2001). In the brains of DYT1 dystonia patients, the distribution of torsinA proteins was the same as in control subjects, without staining of inclusion bodies or the nuclear envelope (Walker et al., 2002a; Rostasy et al., 2003), although inclusion body staining has been reported in brainstem regions (McNaught et al., 2004).

Golgi localization can explain much of the published literature on torsinA localization. First, the torsinA-positive region does not completely overlap with the ER in human brains (Walker et al., 2002a); torsinA staining appears in a small part of cytoplasm (Shashidharan et al., 2000; Walker et al., 2001), indicating that a non-ER compartment was stained. Second, this partial colocalization of torsinA with ER markers needs to be considered carefully, because the optical resolution is limited. In conventional confocal microscopes, the spatial resolution in the lateral (x-y) direction is 200-300 nm, and in the axial (z) direction it is 500-1000 nm (Egner and Hell, 2006; Iwabuchi et al., 2014a), whereas the Golgi and nearby ER are separated by mere tens of nm. Thus, even if the optical signal from the entire Golgi apparatus appears to overlap with the ER (Figs. 5D, 10), this does not support the colocalization of Golgi and ER. Similarly, partial overlap of torsinA signal with the ER signal does not necessarily support the colocalization of torsinA and ER. Therefore the ER staining data do not exclude the localization of torsinA in Golgi. Third, torsinA signal was found in the processes of immature neurons (Granata et al., 2008). This could represent the dendritic Golgi outposts (Horton and Ehlers, 2003; Mitchell et al., 2018) (Fig. 12). Fourth, torsinA was absent from axonal shafts and nerve terminals (Koh et al., 2013), consistent with the lack of Golgi in these regions (Krijnse-Locker et al., 1995; Horton and Ehlers, 2003). Lastly, it is important to note that the focus levels can influence the visible patterns of torsinA staining in somata. They ranged from compact Golgi-like signals to diffuse ER-like signals in the same neurons, in the latter case reflecting out-of-focus stray light from the Golgi above and below the optical section (Figs. 3, 6; widefield and confocal microscopy).

Formally, the torsinA signal localization near Golgi in the current study could be explained by torsinA associating with a subgroup of the ER that segregates from the rest and shows a distribution nearly identical to that of the Golgi apparatus. However, such a structure has not been demonstrated, and, if present, it will still require modification of the existing mis-localization model.

### Implications of the current study

Abnormalities in the Golgi apparatus are associated with neurological disorders (Bexiga and Simpson, 2013; Condon et al., 2013; Neefjes and van der Kant, 2014). Given the preferential localization of torsinA near the Golgi, can the WT- and ΔE-torsinA in this structure affect neuronal functions? Admittedly, the mechanisms whereby torsinA proteins might be retained near the Golgi apparatus (Storrie, 2005; Banfield, 2011) and their function therein are unknown. However, we speculate that torsinA could regulate Golgi-associated functions, including the secretion and intracellular trafficking of proteins. This notion is based on the fact that the Golgi apparatus is an important part of the secretory pathway. Consistent with this hypothesis, torsinA has been reported to regulate the intracellular trafficking of proteins to the plasma membrane. The targets of its activity include the dopamine transporter, norepinephrine transporter, dopamine receptor, adrenergic receptor and ATP-sensitive potassium channel (Torres et al., 2004). In addition, in fibroblasts from DYT1 patients, there was reduced secretion of an artificial reporter (Gaussia luciferase) from ER to plasma membrane (Hewett et al., 2007), although Golgi abnormality was not tested. Moreover, Golgi apparatus is increasingly recognized as organizing centers for microtubules (Efimov et al., 2007; Rivero et al., 2009). This also raises the possibility that mutated torsinA in the Golgi can affect the microtubule-dependent trafficking of important proteins of the secretory pathway, such as synaptic proteins.

In our discussion, we have focused on the relative importance of the Golgi apparatus as a site of torsinA localization. It remains still possible that subtle abnormalities in the nuclear envelope or ER, undetectable at the light microscopy level, underlie the pathophysiology of DYT1 dystonia (Cookson and Clarimon, 2005). However, our study suggests that mutation-associated mis-localization of torsinA might not be a key phenomenon, and warrants reconsideration of how ΔE-torsinA leads to DYT1 dystonia. Discrepancies between the current study and overexpression studies also highlight that the expression level of torsinA dictates its localization. Thus torsinA may belong to a group of modulators whose study at endogenous levels of expression will be critical to understanding the pathophysiology of diseases (Chao and Zoghbi, 2012; Gibson et al., 2013). Our findings also support the notion that rescue strategies based on drastically changing torsinA protein levels could pose potential risks (Martin et al., 2011).

## ACKNOWLEDGEMENTS

The authors thank Drs. Phyllis Hanson (Washington University) for the plasmid construct of WT-torsinA-GFP, Richard Roller (University of Iowa) for the plasmid construct of ΔE-torsinA-GFP, Amy Lee (University of Iowa) for the GRP78/BiP antibody, and Mark Stamnes (University of Iowa) for the GM130 antibody. This work was supported by grants from the Department of Defense (W81XWH-14-1-0301) and the Dystonia Medical Research Foundation (to N.C.H.). The authors declare no conflicts of interest.

